# The genus *Boechera* as a model for apomixis research

**DOI:** 10.1101/2025.01.28.635200

**Authors:** Terezie Mandakova

## Abstract

The genus *Boechera*, a prominent member of the tribe Boechereae within the family Brassicaceae, has emerged as an exceptional model for apomixis research, owing to its unique evolutionary, reproductive, and genomic characteristics. With over 480 genetically distinct taxa and a distribution spanning North America, Greenland, and parts of Asia, *Boechera* exemplifies remarkable ecological and genetic diversity. A hallmark of the genus is its high frequency of gametophytic apomixis, including diplospory and apospory, which, combined with hybridization, drive genetic innovation and environmental adaptability. Moreover, *Boechera* is notable for its diploid expression of apomixis, independent of polyploidy, providing an unparalleled system for dissecting asexual reproduction. Genomic adaptations, such as heterochromatic supernumerary chromosomes, further highlight its evolutionary complexity. Recent advances, including the discovery of critical regulatory genes like *APOLLO* and *UPGRADE2*, have deepened our understanding of the genetic basis of apomixis. These insights position *Boechera* as a cornerstone for elucidating apomictic pathways and leveraging these mechanisms for crop improvement. The genus not only enhances our understanding of apomixis but also offers transformative potential for agriculture.

## The tribe Boechereae (Brassicaceae)

The Brassicaceae family, commonly known as the mustard family, ranks among the largest and most diverse plant families, encompassing nearly 4,000 species across 351 genera [1]. Originating during the late Eocene to Oligocene epochs, this family holds considerable ecological and agricultural significance, thriving across a wide range of ecosystems worldwide [2]. Brassicaceae is classified into two subfamilies: the smaller and less diverse Aethionemoideae, and the expansive Brassicoideae, which comprises five supertribes and 58 tribes, including the tribe Boechereae [2].

The tribe Boechereae stands out for its distinctive reproductive strategies, particularly the extensive presence of apomixis [3]. This tribe encompasses several genera that vary in diversity and geographic distribution. Notably, seven genera - *Anelsonia, Cusickiella, Nevada, Phoenicaulis, Polyctenium, Sandbergia*, and *Yosemitea* - are either mono- or bispecific and predominantly restricted to the western United States. In contrast, the genus *Borodinia* comprises eight species, with seven native to eastern North America and one extending into Siberia and northeastern coastal Asia [4].

The genus *Boechera* is the most diverse and geographically extensive member of the tribe Boechereae. It includes approximately 480 genetically distinct taxa [5], with a distribution that spans from Alaska across much of North America, and extends to Greenland and Siberia for certain species [4]. Recent advancements in phylogenetic analyses utilizing high-resolution genomic tools have significantly enhanced our understanding of the evolutionary relationships within the tribe. These studies have effectively distinguished genera and resolved clades within *Boechera*, providing a robust framework for exploring the evolutionary dynamics and diversification processes of the Boechereae [2, 4].

### Apomixis and reproductive diversity in Boechereae

The tribe Boechereae serves as an evolutionary hotspot for the origin and diversification of apomictic taxa. Within the genus *Boechera*, gametophytic apomixis is observed in over 100 taxa, significantly contributing to genotype and phenotype diversity by stabilizing the genetic outcomes of hybridization and reticulate evolutionary processes [7]. Detailed embryological investigations have identified three distinct mechanisms of gametophytic apomixis in *Boechera*: *Antennaria*-type diplospory, *Taraxacum*-type diplospory, and *Hieracium*-type apospory [7, 8]. Among these, *Antennaria*-type diplospory is notably rare and primarily documented in individuals that predominantly reproduce via *Taraxacum*-type diplospory. *Taraxacum*-type diplospory and apospory, in contrast, are more prevalent and occur at higher frequencies than sexual reproduction in natural populations. Remarkably, both apospory and diplospory are equally represented among *Boechera* taxa, and in seven species, these two modes of apomixis co-occur within the same individual, each expressed at exceptionally high frequencies [7].

Apospory, while extensively studied in *Boechera*, has also been identified in other genera within the tribe Boechereae, including *Borodinia laevigata, Phoenicaulis cheiranthoides* (in its di-, tri-, and tetraploid forms), *Polyctenium fremontii* (tetraploid), and *Sandbergia whitedii* (triploid) [3, 7, 9]. These discoveries highlight the reproductive and evolutionary adaptability across Boechereae, emphasizing its significance as a model system for investigating apomixis in angiosperms.

### Chromosome dynamics and hybridization in *Boechera*

The genus *Boechera* is characterized by a basic chromosome number of *x* = 7, with sexual species predominantly diploid (2*n* = 14). Although primarily self-pollinating (autogamous), interspecific hybridization among sexual diploids frequently results in the formation of allodiploids (2*n* = 14, 15) [3, 10, 11, 12]. These hybrids often adopt apomictic reproduction, a strategy that mitigates sterility typically associated with allodiploidy, thereby enhancing their viability and persistence within natural populations. Remarkably, apomixis in allodiploid *Boechera* - arising from hybridization between sexual diploids -represents a rare phenomenon among flowering plants [7].

A distinctive trait of many apomictic *Boechera* species is their ability to produce unreduced (2*n*) pollen, a phenomenon rarely observed in other apomictic angiosperms [13]. This unique capability allows 2*n* sperm from apomictic diploids to fertilize haploid (1*n*) eggs of sexually reproducing taxa, resulting in the formation of novel triploid apomicts with chromosome numbers of 2*n* = 21 or 22 [5, 12, 14]. Tetraploid apomicts can also emerge through this process, although they occur much less frequently [15, 16]. These apomictic hybrids demonstrate enhanced ecological adaptability compared to their sexual progenitors, enabling them to thrive across diverse environmental niches [17, 18]. However, frequent introgression between sexual and apomictic taxa often obscures morphological and genetic distinctions, complicating the taxonomy of *Boechera* species [5].

### Chromosome structure and evolution

The genomic architecture of the tribe Boechereae has been elucidated through studies on species representing seven genera: *Boechera, Borodinia, Cusickiella, Phoenicaulis, Polyctenium, Nevada*, and *Sandbergia* [3, 5, 9, 19]. These investigations revealed a conserved chromosome base number of *x* = 7 across all analyzed taxa. Comparative analyses with the closely related tribe Halimolobeae, which retains a base number of *x* = 8, suggest that the ancestral genome of Boechereae (*n* = 7) originated via descending dysploidy from an *n* = 8 genome (see **Fig. 1a, b**). This genomic reorganization is estimated to have occurred approximately eight million years ago, coinciding with the divergence of extant Boechereae lineages [3].

**Figure 1.**
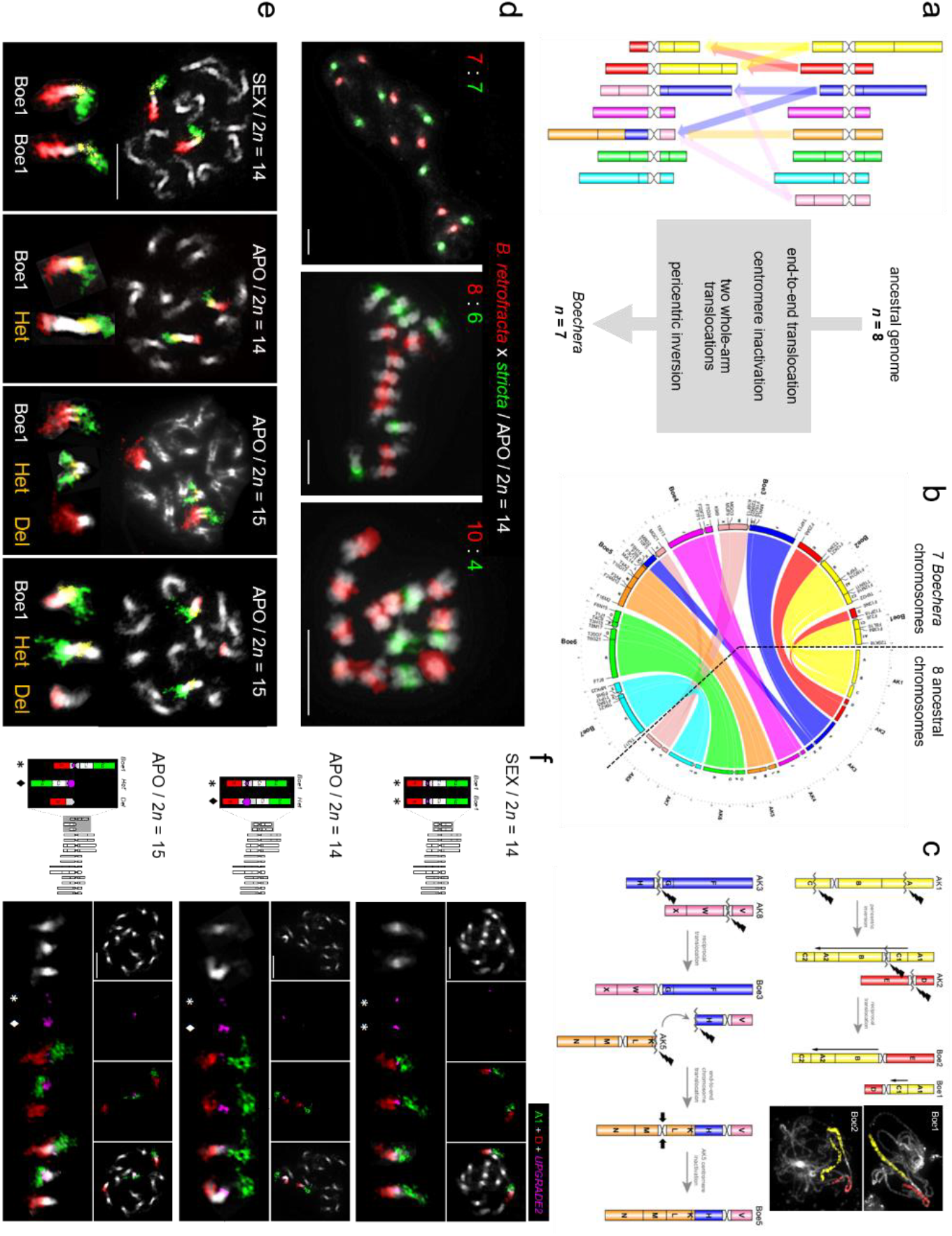
Structure and evolution of *Boechera* genomes and apomixis-associated chromosomes. (a) Origin of the seven *Boechera* chromosomes from an eight-chromosome ancestral genome [modified from 3]. (b) Collinearity between the seven *Boechera* chromosomes and the eight ancestral chromosomes [modified from 19]. (c) Origin of the reshuffled chromosomes Boe1, Boe2, Boe3, and Boe5 [modified from 3]. (d) Genomic *in situ* hybridization showing variable numbers of parental chromosomes in different apomictic *B. retrofracta* x *stricta* lines (unpublished data). (e) Structure of chromosomes Boe1, *Het*, and *Del* revealed by comparative chromosome painting using probes corresponding to genomic blocks A1 (green), C1 (yellow), and D (red) (unpublished data). (f) Fluorescence in situ hybridization of the UPGRADE2 gene [modified from 27]. All chromosomes are counterstained with DAPI. Scale bars, 10 µm.

Of the seven chromosomes present in Boechereae taxa, three (Boe4, Boe6, and Boe7) remain structurally intact and resemble their ancestral configuration. However, the other chromosomes have undergone extensive reshuffling, including an end-to-end translocation, two reciprocal translocations, and a pericentric inversion, culminating in the formation of chromosomes Boe1, Boe2, Boe3, and Boe5 (see **Fig. 1c**) [3, 19]. While Boechereae generally display genomic stability, specific chromosomal rearrangements - particularly inversions - have been linked to diversification within the tribe [3].

One prominent example is a significant pericentric inversion identified in *Boechera stricta*, which has been associated with ecologically important traits and the differentiation of populations [20]. These findings highlight the role of structural genomic changes in reproductive isolation and the speciation processes that have shaped the evolutionary trajectory of the tribe Boechereae.

The genus *Boechera* has garnered significant scientific attention due to its compact genome size (∼250 Mb), its evolutionary proximity to *Arabidopsis thaliana* [2, 21], and the coexistence of diploid sexual and diploid apomictic lineages. Notably, the presence of diploid apomictic lines in *Boechera* highlights that polyploidy is not a prerequisite for the expression of apomixis in this genus. Genomic *in situ* hybridization studies have shown that apomictic *Boechera* are interspecific hybrids, with variability in the number of parental chromosomes resulting from the substitution of homeologous parental chromosomes (see **Fig. 1d**) [10].

A defining feature of apomictic *Boechera* genomes is the presence of heterochromatic supernumerary chromosomes. These additional chromosomes were first documented in aneuploid apomicts with 15 chromosomes as early as 1951 [14]. Subsequent research identified these smaller, heterochromatic chromosomes, referred to as B-like chromosomes, which are absent in sexual diploids but consistently found in apomictic diploids (2*n* = 15) and triploids (2*n* = 22). These B-like chromosomes were hypothesized to carry genetic elements essential for apomixis [22, 23].

Further cytogenetic investigations revealed the ubiquitous presence of a highly heterochromatic chromosome known as *Het* in all diploid apomicts. Additionally, a smaller derivative chromosome, termed *Del* (as presumed a deletion chromosome), was detected in diploid and triploid apomictic aneuploids [10]. Studies on 14-chromosomal apomicts identified the *Het* chromosome as a modified Boe1 homeolog. This chromosome contains genomic blocks A1, C1, and D, characterized by expanded pericentromeric heterochromatin (see **Fig. 1e**) [19]. In 15-chromosomal aneuploid apomicts, the centric fission of the *Het* chromosome resulted in two new derivatives: a telocentric *Het* chromosome (comprising blocks A1 and C1) and a *Del* chromosome (block D) (see **Fig. 1e**) [19]. These fission-derived chromosomes are stably inherited, a process facilitated by the apomictic reproductive mode, which ensures their persistence in populations.

Interestingly, centric fission in *Boechera* represents the only documented case of this chromosomal rearrangement, leading to ascending dysploidy, within the Brassicaceae family. This unique genomic feature further underscores the genus’s value as a model for studying chromosomal evolution in plants.

### Genetic mechanisms underlying apomixis in *Boechera*

The regulation of apomixis in *Boechera* is orchestrated by multiple genetic factors, each associated with distinct components of the reproductive pathway. Prominent among these are the *APOLLO* gene, implicated in female apomeiosis [24], and *UPGRADE2*, which is linked to male apomeiosis [25]. Genome-wide studies investigating gene expression during ovule development have identified significant alterations in regulatory patterns that are characteristic of apomictic reproduction [26]. The functional elements of apomixis - namely apomeiosis, parthenogenesis, and pseudogamy - exhibit variable inheritance patterns within and between *Boechera* species, as demonstrated through segregation and transmission studies [27]. These findings support the hypothesis that apomixis is governed by the interplay of multiple genetic factors rather than a single genetic determinant.

Further insights into the genomic basis of apomixis have emerged from studies on the chromosomal localization of apomixis-associated genes. The *UPGRADE2* gene has been mapped to the Boe1 homologous chromosomes, where an increased copy number has been identified on the *Het* chromosomes in apomictic diploids (2*n* = 14) and aneuploids (2*n* = 15), respectively (see Fig. 1f) [27]. This amplification correlates with the expansion of pericentromeric heterochromatin on the Het and Het′ chromosomes, suggesting a functional link between chromatin structure and gene dosage. The consistent presence of Het chromosomes containing *UPGRADE2* in all apomictic lineages underscores its pivotal role in controlling apomictic reproduction. Collectively, these findings highlight the critical contributions of chromosomal rearrangements and gene copy number variations to the evolution and persistence of apomixis in *Boechera*.

## Conclusion

The genus *Boechera* serves as an unparalleled model for studying apomixis, particularly its expression at the diploid level, independent of the confounding effects of polyploidy. Its diverse mechanisms of gametophytic apomixis, genomic adaptations such as heterochromatic chromosomes, and the identification of pivotal regulatory genes like *APOLLO* and *UPGRADE2* make *Boechera* a critical system for exploring the genetic, evolutionary, and ecological dynamics of asexual reproduction. This genus not only advances our understanding of apomixis but also provides a foundation for translating these insights into practical applications.

## Future directions

Although apomixis in *Boechera* has been the subject of extensive research for decades, significant gaps remain, including the absence of a complete genome sequence for an apomict and a limited understanding of the epigenetic mechanisms underlying this phenomenon. However, recent technological advancements have set the stage for transformative progress. Long-read sequencing technologies, paired with sophisticated algorithms for genome assembly, now enable the generation of telomere-to-telomere genome assemblies, providing unprecedented opportunities to decode the complex genomic architecture of apomicts. Advances in cytogenetics, integrating chromosome painting and immunolocalization, further allow for validation of chromosome-level assemblies and enable detailed investigations into the three-dimensional structure and spatial organization of chromosomes and subgenomes. These tools promise to illuminate the links between hybridization, chromosome reorganization, and the evolution of apomixis.

Future research should prioritize leveraging these breakthroughs to unravel the intricate genetic and regulatory networks governing apomixis, with a focus on the mechanistic roles of key genes. Investigating the environmental and ecological factors that influence apomixis expression will enhance our understanding of its adaptive significance, while detailed studies of heterochromatic chromosomes could shed light on their evolutionary and reproductive functions. Comparative analyses across apomictic systems may uncover conserved pathways, facilitating broader generalizations.

Furthermore, cutting-edge gene-editing tools such as CRISPR/Cas provide an exciting avenue to experimentally reconstruct plant chromosomes and replay evolutionary trajectories, offering unique insights into the interplay between chromosomal dynamics and reproductive strategies. Ultimately, applying synthetic biology and gene-editing technologies to transfer apomictic traits into crop species holds transformative potential for agriculture, paving the way for the development of high-yield, genetically uniform plants. This progress could address critical challenges in global food security and biodiversity conservation, marking a new era for both basic and applied research in plant biology.

## Acknowledgements

This work was supported by the Czech Science Foundation (projects 21-06839S and 24-11371S) and Masaryk University Grant Agency (project MUNI/R/1268/2022).

